# Complex Patterns of Altered White Matter Structural Connectivity within a ‘subjective valuation network’ in Treatment-Resistant Depression

**DOI:** 10.1101/2025.08.31.673378

**Authors:** Benjamin Maas, Rachel Jones, Christinia Johnson, Angela Jiang, Dwayne W. Godwin

## Abstract

**Background:** Treatment-resistant depression (TRD) poses a significant clinical challenge, demanding a deeper understanding of its neurobiological underpinnings to improve therapeutics. We examined white matter microstructure and structural connectivity in TRD, focusing on the ‘subjective valuation network’ (SVN), which captures motivated behavior, reward processing, and emotional regulation circuits commonly altered in depression. This allowed us to identify potential neuroimaging biomarkers associated with treatment resistance.

**Methods:** Diffusion tensor imaging (DTI) data were acquired from a sample of age-and-gender balanced individuals with TRD (n=44; female = 24, male=11, non-binary/unchecked =7) and non-depressed controls (n=42; female=27, male =12, non-binary/unchecked=5). Tract-Based Spatial Statistics were used to compare whole-brain white matter integrity differences between groups. Probabilistic tractography was then used to assess fractional anisotropy (FA) and white matter structural connectivity within the SVN across groups. SVN was defined a priori based on converging functional connectivity studies and included key regions such as the ventromedial prefrontal cortex (vmPFC), anterior cingulate cortex (ACC), ventral striatum, and insula and their connecting white matter tracts. Additionally, correlations between clinical measures of depression severity and cognition and structural features of the network’s white matter fibers (i.e. FA and structural connectivity) were explored.

**Results:** Compared to non-depressed controls, individuals with TRD exhibited reduced FA within the left uncinate fasciculus, left inferior fronto-occipital fasciculus, and anterior cingulum, supporting widespread white matter integrity degradations in TRD. No correlations were found between FA and depression severity, suggesting a more specific association with anhedonic features. Whole-network measures of FA and structural connectivity of the SVN did not differ between groups. However, specific subcircuits’ structural connectivity within the SVN differed between groups; namely, the white matter tracts connecting the insula-vmPFC, striatum-insula, and striatum-vmPFC. Hyperconnectivity emerged for patients with TRD for tracts connecting the insula-vmPFC and striatum-vmPFC region-pairs, with exhibited hypoconnectivity between the striatum-insula. Exploratory analyses for the TRD group indicated the subcircuits with altered structural connectivity within the SVN correlated with depressive severity. This indicates subcircuit network alterations may associate with greater difficulty experiencing pleasure – a core symptom of depression and a potential marker of treatment resistance.

**Conclusions:** This study provides evidence for wide-spread disruption of white matter microstructure and altered structural connectivity within specific subcircuits of the SVN in TRD. These findings point to an intricate pattern of structural hyper-and-hypo-connectivity within the subcircuits of the SVN which may underlie the core symptoms of TRD. The altered structural connectivity within the SVN may contribute to the pathophysiology of TRD, especially concerning the motivational and emotional deficits associated with anhedonia. Future research employing multimodal neuroimaging techniques and longitudinal designs is warranted to further elucidate the functional consequences of these structural abnormalities and their potential as predictive biomarkers for personalized treatment interventions in TRD. Specifically, investigating how these white matter alterations change with successful treatment or targeted interventions aimed at improving anhedonia could inform more effective therapies for this challenging condition.

## Introduction

### Background

Treatment-resistant depression (TRD) is a more severe form of major depressive disorder (MDD) with significantly worse outcomes, including increased health care costs and rates of suicide attempts, and a reduced quality of life.(1,2) In 2021, 8.9 million U.S. adults were classified as medication-treated MDD and 2.8 million as TRD.(3) Of the total $92.7 billion healthcare costs in 2021 associated with MDD, almost half ($43.8 billion) was attributed to TRD.(3) This highlights the fact that despite lower prevalence, treatment costs are estimated to be 40% higher for TRD than medication-treated MDD.(2)

Patients with TRD face higher risks of serious health comorbidities, including severe cardiac and pulmonary diseases and, diabetes mellitus, along with higher rates of relapse and illness burden.(2,4) These elevated health risks and societal impacts highlight the critical need to identify distinct pathological mechanisms and biomarkers to enable differentiation of TRD from medication-treatable MDD, potentially establishing TRD as a distinct neurobiological subtype of MDD.

Patients are typically diagnosed with MDD based on self-reported and/or observed behavioral symptoms alongside family reports and clinical depression assessments.(5) Of the patients diagnosed with MDD, only 30-40% achieve remission after their first antidepressant medication trial.(2,4,5) Furthermore, approximately 50% of patients with MDD do not respond to standard first line drug interventions with significantly decreasing likelihood of success with each successive antidepressant trial.(2,4,5) Patients who do not respond or achieve remission after two or more adequate pharmacological trials are typically considered to have TRD.(1–3,5) While this is the most common definition for TRD, significant clinical uncertainty remains around this definition,(2,3) particularly regarding what constitutes an “adequate” antidepressant trial in terms of duration and medication adherence.(2,3)

No clinical biomarkers currently exist for MDD or TRD. This stresses the need for biological-and-pathological-rooted markers to diagnose MDD, with a particular need to differentiate TRD earlier. Diagnosing TRD earlier would save patients the arduous trial-and-error medication approach that is the standard of care for MDD treatment, which ultimately results in little-to-no symptom alleviation for the TRD sub-group. Additionally, diagnosing TRD earlier would reduce some of the TRD-associated economic and societal costs.

A growing amount of literature points to potential biomarkers and pathological underpinnings for these depressive disorders. Namely, it has been demonstrated that white matter microstructural integrity and/or structural connectivity may underlie or exacerbate depressive symptoms in both MDD and TRD.(2,6–18) Studies have consistently shown both a reduction in white matter integrity as well as altered patterns of structural connectivity in MDD and TRD.(2,9,12)

Diffusion weighted imaging (DWI) is a non-invasive Magnetic Resonance Imaging (MRI) sequence that measures the diffusion of water molecules within brain matter. DWI enables the evaluation of (DWI-related) metrics used as proxies to quantify white matter microstructural health and structural connectivity.(19–23) One key DWI metric, fractional anisotropy (FA), quantifies the degree of anisotropic water diffusion within brain matter and is typically used as a surrogate for white matter microstructural health.(6,9,24) While FA can indicate disruption in white matter connectivity, it should not be interpreted as a direct proxy for anatomical connectivity.(19)

Using DWI, studies have demonstrated, compared to patients with MDD, those with TRD show lower mean FA in the following tracts: body of the corpus callosum, bilateral superior longitudinal fasciculus, forceps major and minor, as well as bilateral effects on the cingulum and inferior longitudinal fasciculus.(2) Other DWI MDD studies point to altered structural connectivity in the default mode network (DMN) and another network whose regions collectively are known to associate with self-referential cognitive and emotional processing tasks.(15,17,18,25) Comparative functional and structural connectivity studies of MDD and TRD reveal complex relationships, suggesting potential compensatory mechanisms and neurobiological signatures unique to TRD.(2,12,15,16,26–32)

MDD and TRD are both characterized by altered brain network connectivity and dysfunctional processing of subjective feelings, particularly within brain networks associated with motivated decision-making behavior, momentary subjective valuation, and external reward valuation.(2,14–18,25–38) Converging functional imaging evidence identifies four core hubs, or regions of interests (ROIs), associated with momentary subjective valuation during motivated decision-making behavior: the anterior cingulate cortex (ACC), ventral-medial prefrontal cortex (vmPFC), ventral striatum, and insula.(29,35,36) These ROIs are crucial for computing and updating reward values, motivation, and salience signals while engaged in value based decision making.(29,35,36,39–42) For these reasons, we operationally define these ROIs here as a putative ‘subjective valuation network’(SVN) reflecting their well-established roles of processing subjective valuations throughout motivated decision-making behavior within the broader reward and salience systems. Our analysis focused on six ROI-ROI pairs known to connect via established white matter tracts in the SVN that are specifically associated with motivated-decision-making behavior: (i) striatum-vmPFC, (ii) striatum-insula, (iii) vmPFC-ACC, (iv) insula-ACC, (v) insula-vmPFC, and (vi) ACC-striatum.

Recent literature links TRD to aberrant emotional processing and momentary subjective valuation, motivational deficits, blunted reward responses, and anhedonia.(2,28–31,36) MDD studies have demonstrated functional hypoconnectivity within the DMN and decreased functional switching between executive control and DMN networks normally mediated by the salience network (SN).(2,12,17,18,25,27,29,31–34,36,37) Normally, network switching is triggered by the ACC and insula which help detect salient internal and external events.(14,25,32,33,43) These functional aberrations within the DMN and SN are theorized to underlie the excessive rumination behaviorally observed in TRD and may contribute to a maladaptive focus on internal states.(2,12,17,18,25,27,29,31–34,36,37,43)

Recent scientific findings motivate our investigation of potential white matter differences of the SVN in TRD. (2,12,14–18,25–28,36,42,44) The National Institute of Mental Health’s Research Domain Criteria (RDoC) provides a framework for investigating psychiatric illness subtypes that acknowledges the heterogeneity in underlying mechanisms within a categorized disease state. Building on this approach, we investigate white matter alterations in structural integrity and connectivity within the SVN, and its pathological relevance to TRD as a distinct subtype of depression.(12,27,29–31,36)

### Study Objectives

In this study we aim to compare white matter structural abnormalities (using both FA [(micro)structural integrity] and probabilistic structural connectivity) in TRD vs. controls, focusing on the SVN. Specifically, we examine:

1) Whole-brain major white matter tract differences in FA;
2) Network FA differences of the SVN;
3) Structural connectivity differences, examining both:
  a. Composite network connectivity across all six ROI-ROI pairs of interest;
  b. Individual ROI-ROI structural connectivity between SVN ROIs: striatum-vmPFC, striatum-insula, vmPFC-ACC, insula-ACC, insula-vmPFC, and ACC-striatum;
4) Relationships between clinical measures, age, and white matter characteristics (whole-brain FA, network FA, and ROI-ROI tract-specific SVN structural connectivity)

The clinical significance of this study is to further understand possible white matter differences underlying anhedonia, reward hyposensitivity, and persistent depressive symptomology in TRD. Particularly, both at macro-network-and micro-subcircuit levels of the SVN, offering insight into potential pathological distinctions and neuroimaging biomarkers unique to TRD.

## Material and Methods

### Participants

Participants were recruited and provided written informed consent under Wake Forest University School of Medicine IRB00056131, which corresponds to a larger study under which this present study is nested. The larger longitudinal Electroconvulsive study (ECT) enrolled participants into three groups: (1) a control group with self-reported history of no diagnosis or treatment for depression (2) a MDD group with self-reported history of diagnosis and current treatment for depression but never receiving, seeking, or being offered ECT and (3) a pre-ECT group enrolled to receive a full course of ECT treatments at Atrium Health Wake Forest Baptist Outpatient Psychiatry and Behavioral Health. Exclusion criteria for all participants included contraindication for MRI.

All neuroimaging and clinical data were collected at baseline, prior to the administration of any ECT interventions.

#### Study Groups

An age-and-gender balanced sample of 86 participants underwent structural MRI and DWI. This study consisted of non-depressed control (n = 44) and TRD (n = 42) groups. The control group reported no history of or treatment for depression. The TRD group included participants who were in current treatment for depression with self-reported history of two or more trials of pharmacological interventions, the common clinical definition for TRD, as well as participants already clinically diagnosed with TRD and seeking second-line treatments (in this study, ECT). (5) Rather than separating participants by treatment groups as in the parent study, by combining these subgroups, these inclusion criteria in the present study capture the spectrum of TRD severity; from those unlikely to achieve remission through pharmacological interventions having attempted 2 or more medications to those seeking second-line treatments. This allowed for the investigation of neural signatures across the disorder’s continuum that are potentially unique to and shared across all TRD states.(5,45)

### Data Collection

#### Imaging Acquisition

High-resolution multi-shell DWI data and T1 structural images were collected using a Siemens MAGNETOM 3Tesla Skyra whole body scanner using the following protocols. The DWI data was acquired with simultaneous multi-slice acquisition; isotropic 2×2×2 mm^3^ resolution; TR = 3500 ms; TE 107 ms; FOV 25.6 cm. The DWI data included four non-diffusion weighted (b = 0) volumes and three sets of 60 diffusion-weighted volumes at three increasing gradients (b-values) of b =1, 2, & 3 x 10^3^ s/mm^2^, with gradient orientations isotopically distributed. T1-weighted structural images were also collected for anatomical reference, use in image registrations and normalizations, and DWI data preprocessing.

#### Neuropsychological evaluations, clinical measures, and demographical data

Three neuropsychological evaluations were collected for all study participants. Clinical depression severity was assessed using the Hamilton Rating Scale for Depression (HAMD) and the Patient Health Questionnaire-9 (PHQ9).(46) Cognitive status was evaluated using the Montreal Cognitive Assessment (MOCA).(47) Age and gender demographics were also collected.

### Imaging Data Processing and Connectivity Measures

#### Preprocessing

Preprocessing and analysis protocols were implemented using the FMRIB Software Library (FSL) neuroimaging software.(48) Briefly, this included estimating of and correcting for both susceptibility induced distortions and motion artifacts, removing extraneous non-brain tissue from images (e.g. skull, scalp, etc.), and correcting for induced eddy currents during DWI acquisition.(48)

#### Tensor Model Fitting

The diffusion tensor model was fit for each subject using a single-tensor model at each voxel, creating one dominant fiber orientation per voxel. During this process, FA, mean diffusivity, and the first through third eigenvector and eigenvalue images (often referred to as maps) were produced. The FA images were subsequently used in downstream global metric analyses (i.e. Tract Based Spatial Statistics (TBSS) and FA network SVN analyses).(48)

#### Data preparation for probabilistic tractography and connectivity

For probabilistic tractography, Bayesian estimation of the diffusion parameters was obtained using a “ball-and-stick” model at each voxel and Markov Chain Monte Carlo sampling was used to capture their uncertainty. This approach provided the ability to produce robust estimations of probabilistic maps of white matter connections, rather than deterministic ones. The outputs from these analyses were used as inputs for probabilistic tractography to create probabilistic maps of white matter connections of the SVN for each subject.(22,23)

Detailed FSL preprocessing and probabilistic tractography commands are provided in the Supplementary Methods.

#### Tract Based Spatial Statistics (TBSS)

TBSS was used to localize group FA differences in whole-brain white matter tracts.(49) TBSS involves a robust voxelwise non-parametric permutation testing pipeline. This included non-linearly registering each subject’s FA maps to the 1 mm^3^ standard Montreal Neurological Institute (MNI152) coordinate space via the use of FMRIB58_FA as an intermediate template image.(50,49) A skeletonized study-specific cohort-based mean skeleton was created and used for all subsequent analyses and contrasts implemented.

#### TBSS Contrasts Conducted

Design matrices were created to perform the following contrasts to test for significant differences in regions (or cluster of voxels) in whole-brain white matter tracts between groups: (1) FA values higher in the Control vs. TRD group (i.e. FA; Control > TRD). (2) FA values higher in the TRD vs. Control group (i.e. FA; TRD > Control). (3) FA values higher in the Control vs. TRD group while covarying for age (i.e. FA; Control > TRD with age as a covariate) as FA is known to negatively correlate with age.(51) Age was z-scored within the design matrices.

Additional TBSS contrasts were designed to test if HAMD, PHQ9, and MOCA evaluations significantly correlated with FA values in whole brain major white matter tracts within the design contrast of FA; Control > TRD with age as a covariate as well as independently of group membership. Age and clinical measures were z-scored to ensure comparability across the differently scaled measures as well as to promote numerical stability of the model.

#### Cluster Identification and Anatomical Labeling

Voxelwise non-parametric permutation group label shuffling was set at 500 permutations to account for multiple comparisons via FSL’s *randomise* tool.(49) Statistical maps generated by *randomise* used threshold-free cluster enhancement (TFCE) with an alpha level of p < 0.05.

Significant clusters were extracted using FSL’s *cluster* tool. To assign anatomical labels, the binarized masks were queried against the JHU ICBM-DTI-81 white matter tractography atlas using FSL’s *atlasquery* tool. This enabled localization of clusters to major white matter tracts based on peak voxel coordinates and tract probability overlap. (Detailed command-line procedures are provided in the Supplementary Methods).

#### Defining the ‘subjective valuation network’ (SVN)

Four bilateral anatomical ROI masks of the ACC, vmPFC, insula, and ventral striatum were delineated using FSL constituting the SVN.(48) The insula was retrieved from the MNI Structural Atlas, ventral striatum from the Oxford Imanova Striatal Connectivity Atlas, the vmPFC from https://identifiers.org/neurovault.image:396585, and the ACC from https://identifiers.org/neurovault.image:11988. (50,52) All anatomical ROI images were binarized to create binary masks. The images (vmPFC and ACC) retrieved from neurovault.org were resampled to MNI152 1 mm^3^ standard space. The insula and ventral striatum masks created via FSL were also in MNI152 1 mm^3^ standard template space.

#### Probabilistic Tractography

In FSL, via *ProbtrackX,* probabilistic tractography was conducted in network mode (set at n = 1000 seeds/voxel) using the masks created for the SVN for each subject, producing voxelwise probabilities of white matter fiber connections. The non-linear transformation warp files created during TBSS’s robust registration pipeline from each subject’s diffusion space to standard MNI152 1 mm^3^ template space was supplied via *ProbtrackX’s xfm* option. Inverse non-linear transformation files (from subject diffusion space to MNI 1 mm^3^ template space) were created via FSL’s tool *invwarp* and supplied via *ProbtrackX’s invxf* option. This ensured rigorous anatomical alignment of the SVN ROIs to each subject’s diffusion space.

This analysis yielded two main files for each subject: (1) the probabilistic map of white matter connections image file (in MNI152 1 mm^3^ template space) with probabilistic streamline counts in each voxel of the number of white matter fibers connecting all ROIs in the SVN and (2) a connectivity matrix with streamline counts of plausible white matter fiber connections for each ROI-ROI pair in the SVN (as a text file with an associated waypoint vector text file denoting the number of total streamlines sent out from each ROI respectively).

#### Mean FA values of the SVN

To calculate the mean FA of the SVN and minimize false positives consistent with previous heuristics and methodological studies in the tractography literature, a stringent threshold of 90% was applied to each subjects’ generated image of probabilistic white matter connections SVN map.(20,21,23,53) This resulted in only the top 10% of probabilistic streamline counts of white matter connections being retained in the SVN image. Each image was binarized and multiplied by each subject’s respective FA image that was normalized to the MNI 152 1 mm^3^ space (through TBSS’s robust normalization pipeline). Non-zero mean values for each image were outputted with a minimum threshold value of 0.15 (FA < 0.15 is typically considered non-white brain matter) obtaining each subject’s mean FA value for the SVN.(13,51) Figure 1 below visualizes and summarizes the pipeline implemented to calculate the mean FA values of the SVN.

**Figure 1:**
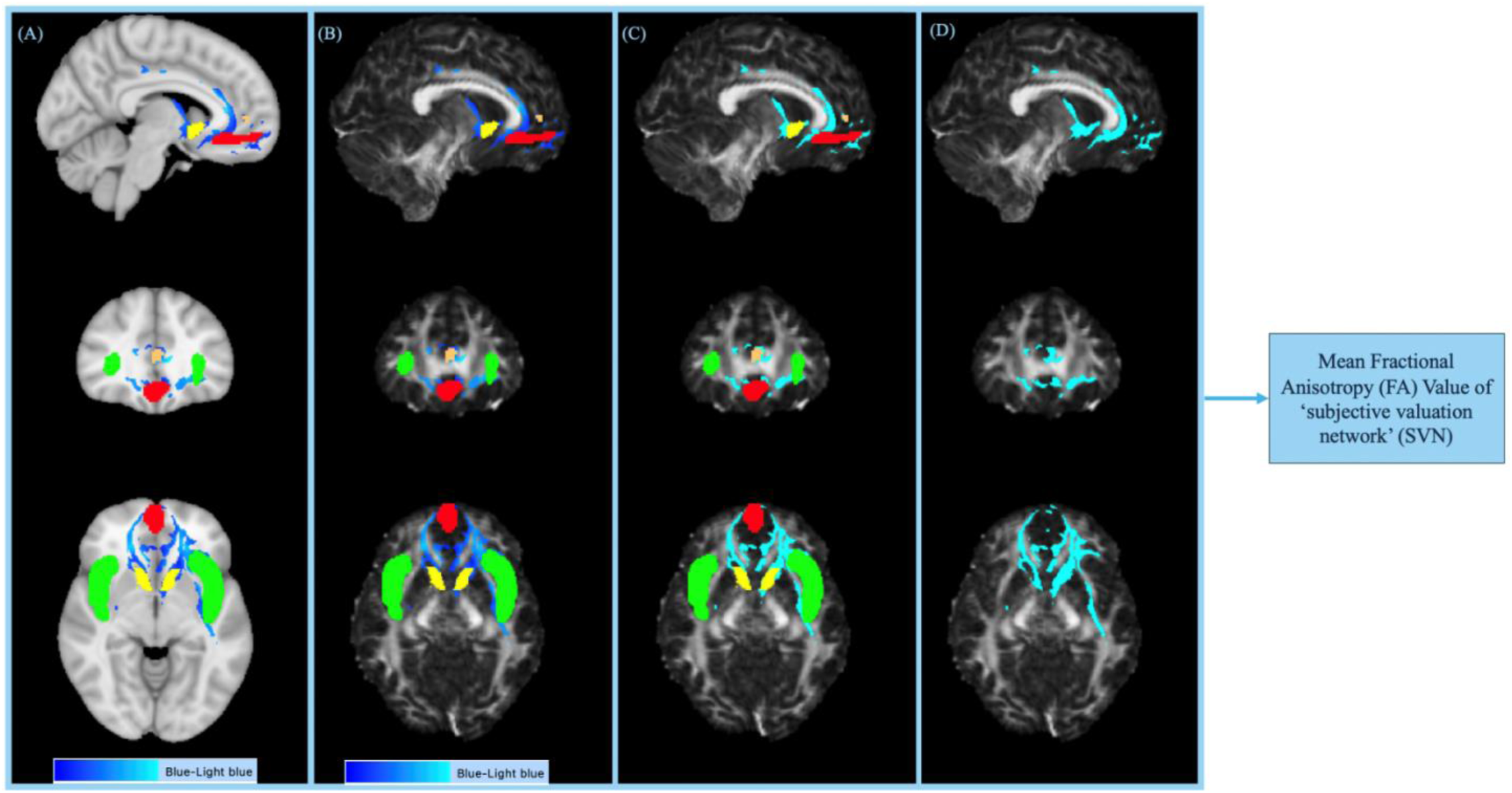
Probabilistic Tractography & Mean Fractional Anisotropy of the ‘subjective valuation network’ (SVN): Probabilistic tractography of the SVN was conducted producing probabilistic maps of white matter connections of the SVN. Depicted in Figure 1(A) and 1(B) is an example subject’s probabilistic tractography output thresholded for the top 10% of streamline counts overlaid the MNI152 1mm^3^ template standard brain and subject’s FA map in MNI152 mm^3^ space respectively, with the binary anatomical masks of the SVN also overlaid; in figure 1(A) and (B), probabilistic tractography is shown on a Blue-Light Blue scale with increasing streamline probability. Throughout Figure 1, masks of the SVN regions are depicted in the following colors: Anterior cingulate cortex (copper), striatum (yellow) ventro-medial prefrontal cortex (red), and insula (green). Figure 1(C) displays the probabilistic tractography output of the SVN post-binarized overlayed the subject’s respective FA map, with the SVN binary SVN masks included. Figure 1(D) displays the probabilistic tractography post-binarized overlaid the subject’s respective FA map, without any of the SVN masks overlayed. Figure 1(D) provides insight into how the mean FA value of the SVN is produced; by multiplying the subject’s binarized probabilistic tractography of the SVN by the subject’s respective FA map. (Detailed command line procedures are included in the supplementary methods).

Detailed command-line procedures are provided in the Supplementary Methods for BedpostX, ProbtrackX, and the assessment of mean values of the SVN.

#### Individual ROI-ROI Structural Connectivity of the ‘subjective valuation network’

Structural connectivity measures were produced for each ROI-ROI pair of interest within the SVN using the produced connectivity matrices [(i) striatum-vmPFC, (ii) striatum-insula, (iii) vmPFC-ACC, (iv) insula-ACC, (v) insula-vmPFC, and (vi) ACC-striatum]. The structural connectivity index is used to estimate the mutual structural connectivity of white matter between two ROI pairs in any connectivity matrix produced through *ProbtrackX* conducted in network mode; here, done for the six pairs of ROI-ROI connections of interest in the SVN.

Each subject’s connectivity matrix consists of a ROI x ROI streamline count for each respective pair with 4 ROIs, a 4 x 4 matrix is produced. Additionally, a 4 x 1 waytotal vector is produced (corresponding to each row of the connectivity matrix and total streamline count sent out per ROI). The waytotal values were used to normalize the connectivity matrix to normalize the structural connectivity measure. Included in the supplementary material (Figure S1) displays the structure of a generic connectivity matrix and waypoint vector of the SVN, and additionally includes the equations used to calculate forward, reverse, and structural connectivity values retrieved from FSL guidelines which were adapted into python code to batch process the assessed metrics.

#### Composite Network Connectivity

In addition to the individual structural connectivity measures assessed, an overall aggregate mean network connectivity score for each subject was calculated. This was done as a proxy to test for overall differences in network structural connectivity across groups. First, the mean value for each subject’s 6 ROI-ROI pairs and connections of interest were calculated respectively, creating a measure of composite network connectivity for each subject. Next, group-level ‘*composite network connectivity’* scores were calculated by averaging individual scores within the control and TRD groups.

#### Statistical Analyses

All statistical analyses were performed in RStudio (version 2024.12.0+467) to test for significant differences between the control and TRD groups in each of the computed metrics: (1) mean FA of the SVN, (2) structural connectivity measures for each individual ROI-ROI pair of interest in the SVN (3) the ‘*composite network connectivity’* score.

Normality was assessed separately for the control and TRD groups for each metric of interest (Shapiro–Wilk, p>0.05). Parametric tests (two-sample t-test) were applied only when both groups met normality criteria; otherwise, non-parametric Mann–Whitney U tests were used.

Multiple comparisons of the structural connectivity across the six ROI–ROI pairs were controlled via False Discover Rate (FDR). Results were considered significant at p<0.05 (FDR-corrected)

#### Correlational analyses of the mean FA value of the SVN

For the mean FA value of the SVN we assessed correlations with HAMD, PHQ9, and MOCA both in relation to control (n = 44) and TRD (n = 42) groups separately, as well as independently of group membership with all participants (n=86). Prior to correlational analyses age and clinical measures were z-scored to ensure comparability across variables between the different scales allowing for standardized effect size interpretation. Normality checks were repeated within each analysis streamline before applying parametric or nonparametric tests. Shapiro-Wilk test (p>0.05) was used to test for normality prior to conducting each correlational analysis. Pearson correlations were used if data met normality criteria; otherwise, Spearman’s rank correlation was applied. If age significantly correlated (p<0.05) with the FA value of either group, or with the FA value independent of group membership, age was included as a covariate in partial correlation analyses. We applied an FDR correction across correlational tests and results were considered significant at p<0.05 (FDR-corrected).

#### Additional exploratory analyses for significantly different ROI-ROI pairs

We planned to explore the relationship between age and clinical measures, independent of group membership, of each ROI-ROI pair of the SVN found to significantly differ between groups (Control vs. TRD) to highlight potential relationships for future hypothesis-driven research; this exploratory assessment was done with the motivation of offering potential further explanation and insight into any findings of hypo-or-hyper subcircuit structural connectivity.

First, normality was assessed for each ROI-ROI pair’s structural connectivity using Shapiro-Wilks tests (p>0.05). If the structural connectivity measure met normality, Pearsons correlational tests were applied, otherwise Spearman’s rank correlational tests were used. If age was found to significantly correlate with the structural connectivity of an ROI-ROI pair (p<0.05), age was used as a covariate in the correlational analyses with each respective clinical measure (HAMD, PHQ9, MOCA). Given the targeted nature of these exploratory analyses, we elected to report uncorrected p-values while offering insight into potential directional associations with clinical measures for targeted future research work.

## Results

### Sample Demographics

The sample comprised of 86 participants (Control: n = 44, TRD: n = 42) balanced in both age (Mann-Whitney U Test; p>0.05) and gender (*x^2^*= 0.71). The TRD group exhibited higher depression severity and lower MOCA ratings. The TRD subjects had a mean HAMD score of 26.40, indicating very severe depression (HAMD scores ≥ 23). The TRD subjects also had a mean PHQ9 score of 16.08, indicating moderately severe depression (PHQ9: 15-19). Table 1 presents demographic and clinical measures in the control and TRD groups, along with the statistical tests and p-values used to assess between-group differences.

**Table 1.** Group Demographic and Clinical Measures: Descriptive Statistics, Between-Group Comparisons, and Statistical Tests: Summary of age, gender, and clinical measures (HAMD, PHQ-9, MOCA) for the control and the TRD groups. Group comparisons were conducted using Mann-Whitney U Tests for continuous variables and a chi-square test for gender.

| Table 1: Group Demographic and Clinical Measures: Descriptive Statistics, Between-Group Comparisons, and Statistical Tests |  |  |  |  |
| --- | --- | --- | --- | --- |
|  | Control | TRD | Test | P-Value |
| Age Mean | 40.39 | 41.25 | Mann-Whitney U Test | 0.84 |
| Age SD | 15.09 | 15.80 | (Not Applicable) | (Not Applicable) |
| Age median | 40 | 41 | (Not Applicable) | (Not Applicable) |
| Age range | 19-66 | 19-69 | (Not Applicable) | (Not Applicable) |
| Gender | Female = 27<br>Male = 12<br>Non-Binary/Unchecked =5 | Female = 24<br>Male = 11<br>Non-Binary/Unchecked=7 | Chi-Square | 0.71 |
| HAMD Mean | 1.91 | 26.40 | Mann-Whitney U Test | < 0.001 |
| HAMD SD | 2.10 | 3.00 | (Not Applicable) | (Not Applicable) |
| HAMD Median | 1 | 27 | (Not Applicable) | (Not Applicable) |
| HAMD Range | 0-7 | 18-30 | (Not Applicable) | (Not Applicable) |
| PHQ9 Mean | 1.66 | 16.08 | Mann-Whitney U Test | < 0.001 |
| PHQ9 SD | 2.48 | 6.95 | (Not Applicable) | (Not Applicable) |
| PHQ9 Median | 1 | 16 | (Not Applicable) | (Not Applicable) |
| PHQ9 Range | 0-13 | 1-30 | (Not Applicable) | (Not Applicable) |
| MOCA Mean | 27.14 | 18.75 | Mann-Whitney U Test | < 0.001 |
| MOCA SD | 2.05 | 9.10 | (Not Applicable) | (Not Applicable) |
| MOCA Median | 27.5 | 19.5 | (Not Applicable) | (Not Applicable) |
| MOCA Range | 20-30 | 0-37 | (Not Applicable) | (Not Applicable) |

### Voxelwise TBSS analysis

Voxelwise whole-brain TBSS analysis revealed significantly reduced FA clusters in TRD (Control > TRD) within several major white matter tracts, with no significant clusters found for TRD > Control. The significant clusters found for the Control > TRD contrast were located within the following white matter tracts; forceps minor (anterior corpus callosum), splenium (posterior corpus callosum), inferior fronto-occipital fasciculus (IFOF), and inferior longitudinal Fasciculus (ILF). Table 2 provides summary statistics for the peak voxels and anatomical locations of significant white matter clusters where FA was lower in TRD compared to controls (Control > TRD contrast).

**Table 2.** TBSS (Fractional Anisotropy) Significant Clusters: Control > TRD Contrast. Peak voxel coordinates and cluster characteristics for white matter regions showing significantly reduced FA in the TRD group comparted to controls. Results were obtained using FSL’s *randomise* (500 permutations, TFCE-corrected at p < 0.05). Anatomical labels were determined by querying the JHU ICBM-DTI-81 white matter tractography atlas.

| Cluster Index | Tract | Vox els | 1-p-MAX | 1-p-MAX X (mm) | 1-p-MAX Y (mm) | 1-p-MAX Z (mm) | 1-p-COG X (mm) | 1-p-COG Y (mm) | 1-p-COG Z (mm) |
| --- | --- | --- | --- | --- | --- | --- | --- | --- | --- |
| 7 | Forceps Minor | 752 | 0.958 | 8 | 26 | -4 | 6.42 | 24.5 | 13.2 |
| 6 | Splenium of corpus callosum | 597 | 0.958 | -11 | -41 | 21 | -14.3 | -38.5 | 24.6 |
| 5 | Splenium of corpus callosum | 520 | 0.958 | 24 | -45 | 25 | 17.2 | -35.3 | 30.1 |
| 4 | Splenium of corpus callosum | 97 | 0.95 | 22 | -48 | 9 | 22.3 | -49.3 | 14.3 |
| 3 | L inferior fronto-occipital fasciculus & L longitudinal Fasciculus | 92 | 0.95 | -28 | -70 | 17 | -29.6 | -67 | 21.4 |
| 2 | L inferior longitudinal fasciculus | 24 | 0.95 | -26 | -79 | 2 | -27 | -77.4 | 3.96 |
| 1 | L inferior fronto-occipital fasciculus | 1 | 0.95 | -27 | -78 | 8 | -27 | -78 | 8 |

Abbreviations: Coordinates (mm) = millimeter; COG = Center of Gravity; L = Left; Max = Maximum; 1 – P = Significance level = 1 – 0.95 = P < 0.05

### Age-Adjusted Analysis

In the TBSS (Control > TRD) contrast with age (z-scored) as a covariate, the same fronto-callosal clusters remained significant indicating these differences are independent of age effects. The persistent presence of significant voxels in these fronto-callosal clusters indicates robust group differences and reduced FA in TRD within these clusters and white matter tracts. Table 3 summarizes the TBSS results for the Control > TRD contrast with age included as a covariate.

**Table 3.** TBSS Significant Clusters: Control > TRD Contrast (Age Adjusted) Peak voxel coordinates and cluster characteristics for white matter regions showing significantly reduced FA in the TRD group comparted to controls with age (z-scored) included as a covariate. Cluster locations and peak voxel coordinates are reported. Results were obtained using FSL’s *randomise* (500 permutations, TFCE-corrected at p < 0.05). Anatomical labels were determined by querying the JHU ICBM-DTI-81 white matter tractography atlas.

| Cluster Index | Tract | Voxels | 1-p MAX | 1-p-MAX X (mm) | 1-p-MAX Y (mm) | 1-p-MAX Z (mm) | 1-p-COG X (mm) | 1-p-COG Y (mm) | 1-p-COG Z (mm) |
| --- | --- | --- | --- | --- | --- | --- | --- | --- | --- |
| 3 | Splenium of corpus callosum | 2867 | 0.958 | -11 | -40 | 21 | -7.45 | -47.2 | 21 |
| 2 | Forceps minor | 682 | 0.956 | 10 | 33 | 2 | 6.83 | 24.9 | 12.9 |
| 1 | Forceps minor | 8 | 0.95 | -37 | -41 | 18 | -36.6 | -40.1 | 19 |

Abbreviations: Coordinates (mm) = millimeter; COG = Center of Gravity; L = Left; Max = Maximum; 1 – P = Significance level = $1 - 0.95 = P < 0.05$
Abbreviations: Coordinates (mm) = millimeter; COG = Center of Gravity; L = Left; Max = Maximum; $1 - P = \text{Significance level} = 1 - 0.95 = P < 0.05$

### Non-significant Clinical Correlations

Neither HAMD, PHQ9, nor MOCA scores showed significant voxelwise correlations with FA at p < 0.05 (corrected).

These findings of disruptions in interhemispheric white matter integrity (forceps minor) and posterior callosal fibers (splenium) are consistent with other literature on TRD TBSS and white matter microstructural integrity studies.(2,6–14,16–18,24,26,54) The absence of correlations between TBSS-derived FA reductions and clinical severity measures, suggests that differences in integrity in global whole-brain white matter tracts may miss more focal microstructural changes, potentially contained within the SVN that may be uniquely pertinent to TRD pathology.

## Macro-Level Indices of the SVN

### Mean FA value of the SVN

There were no significant group differences in mean FA values of the SVN (t-test, p = 0.78). Multiple comparison correction was not applicable as this statistical test involved comparison of one measure (i.e. mean FA value of SVN). This result suggests that FA alone may not sufficiently capture white matter differences within the SVN. Figure 3(A) presents violin plots illustrating the distribution of mean FA values of the SVN across the control and TRD group.

## Correlational analysis of mean FA of the SVN

### Control Group

Additionally, age and clinical measures (HAMD, PHQ9, MOCA) did not significantly correlate with the mean FA of the SVN within the Control group (Age: p = 0.087 FDR-corrected; HAMD: p = 0.28 FDR-corrected; PHQ9: p = 0.97 FDR-corrected; MOCA: p = 0.97 FDR-corrected).

### TRD Group

However, the mean FA value of the SVN did significantly correlate with age within the TRD group (Age: p = 0.0060 FDR-corrected). The mean FA of the SVN for the TRD group did not significantly associate with the assessed clinical measures (HAMD: p = 0.50 FDR-corrected; PHQ9: p = 0.97 FDR-corrected; MOCA: p = 0.93 FDR-corrected). Age had a modest to moderate negative correlational effect size associated with the mean FA value of the SVN (ρ = –0.45). Figure 2 displays the association between age and mean FA of the SVN shown separately for the control and TRD groups.

**Figure 2:**
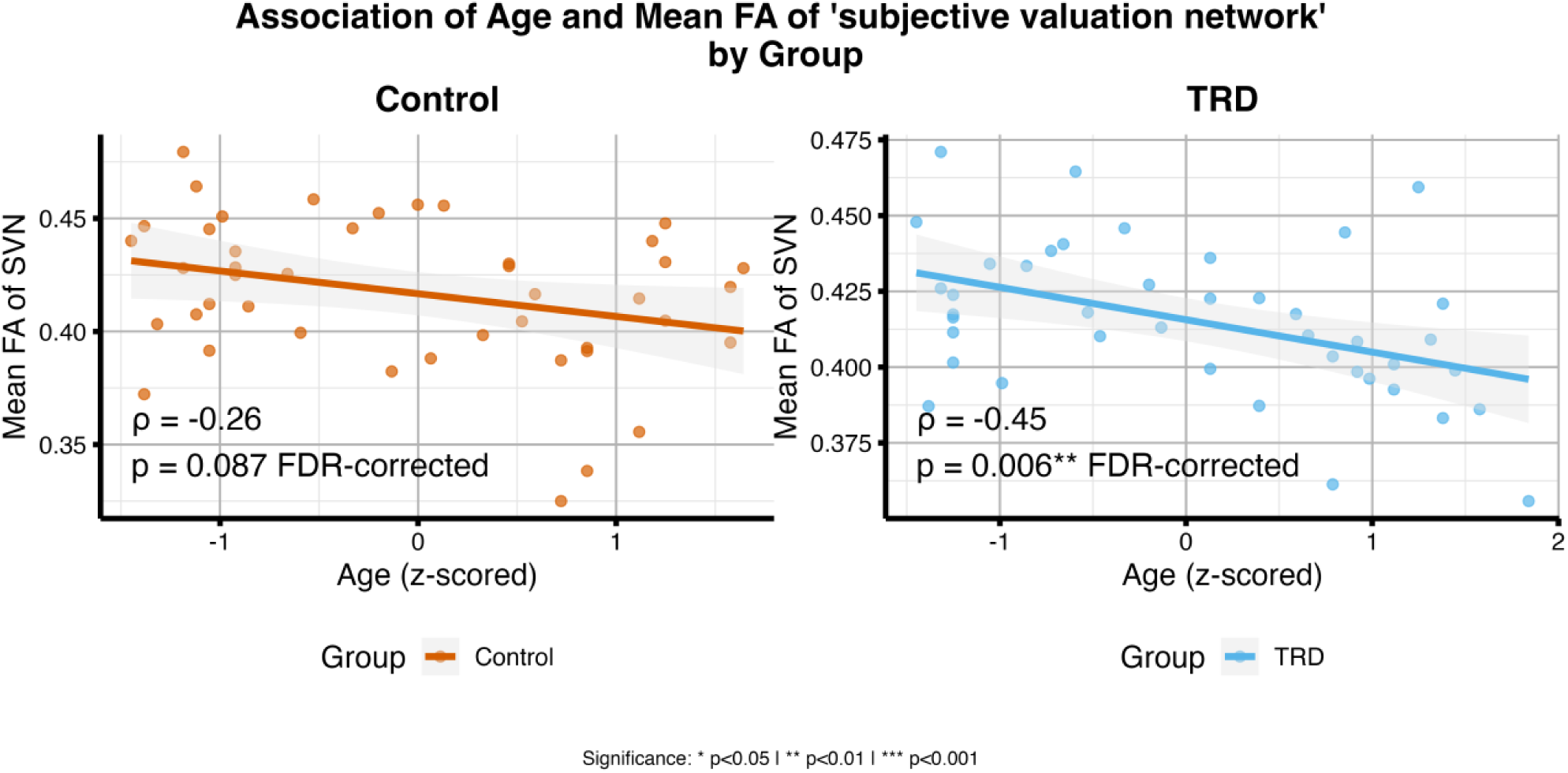
Association between Age and mean Fractional Anisotropy (FA) of the ‘subjective valuation network’ (SVN) by group. Scatterplots with fitted regression lines show the relationship between age (z-scored) and mean FA of the SVN in the control (left) and TRD (right) groups. A significant negative correlation was observed in the TRD group (ρ = –0.45; p = 0.0167 FDR-corrected), while no significant association was found in the control group (ρ = –0.26; p = 0.2796 FDR-corrected). Shaded areas represent 95% confidence intervals. Individual data points are overlaid.

**Figure 3:**
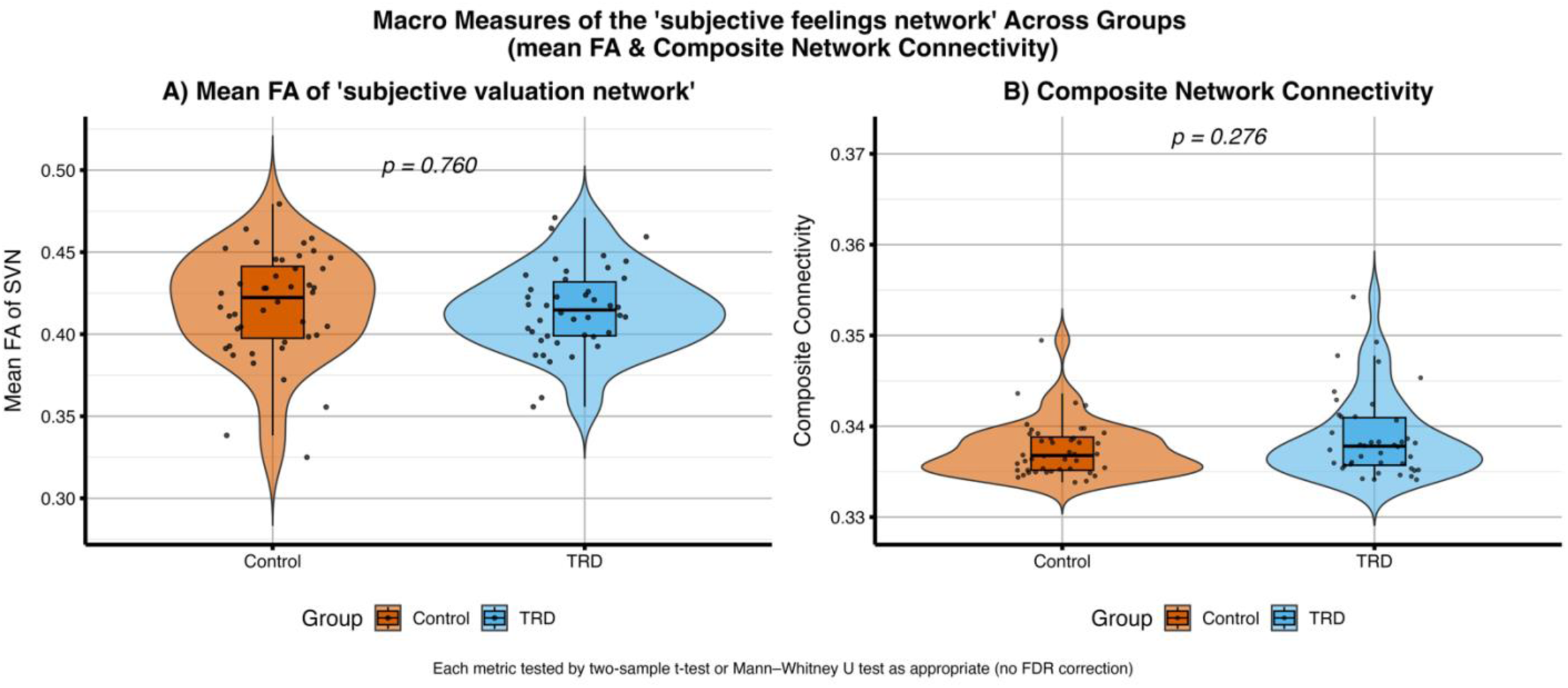
Macro-level white-matter measures of the ‘subjective valuation network’ (SVN) across control and TRD groups. (A) Mean fractional (FA) of the SVN did not differ between Control and TRD participants (two sample t-test, p = 0.760). (B) *Composite Network Connectivity* of the SVN also showed no group differences (Mann-Whitney U Test, P = 0.276). In both panels, violin shapes depict the full distribution with embedded boxplots indicating median (+/−) inter quartile ranges, and individual data points overlaid.

### Independent of Group Membership

A similar pattern emerged for the independent of group membership correlational analyses of the mean FA of the SVN with age and the clinical measures: the mean FA of the SVN significantly correlated with age (Age: p = 0.02 FDR-corrected, ρ = –0.32) but not with clinical measures (HAMD: p = 0.97 FDR-corrected; PHQ9: p = 0.97 FDR-corrected; MOCA: p = 0.91 FDR-corrected).

### Composite network connectivity

Similarly, the *composite network connectivity* did not significantly differ between the Control and TRD groups (Mann-Whitney U; p = 0.28). Similarly, to the mean FA of the SVN two sample t-test, no FDR correction was applied to the single Mann-Whitney U-test. Figure 3(B) displays a violin plot comparison of the *‘composite network connectivity’* measure across groups.

Macro-level measures of the SVN did not distinguish TRD from controls, implying that any white matter abnormalities may manifest at more localized, subcircuit levels within the network.

### Structural connectivity of individual tracts within the SVN

Finally, we assessed ROI-ROI connectivity from the *Probtrackx* connectivity matrix (i.e. individual structural white matter connectivity measures), identifying hyper-and-hypo structurally connected tracts in TRD.

Significant group differences were found in the following connections:

- **(a) insula-vmPFC:** *TRD > Control* (p = 0.0045 FDR-corrected)
- **(b) striatum-insula:** *Control > TRD* (p = 0.0015 FDR-corrected)
- **(c) striatum-vmPFC:** *TRD > Control* (p = 0.040 FDR-corrected)

Figure 4 displays the specific white matter tracts found to be significantly different between the control and TRD groups.

**Figure 4:**
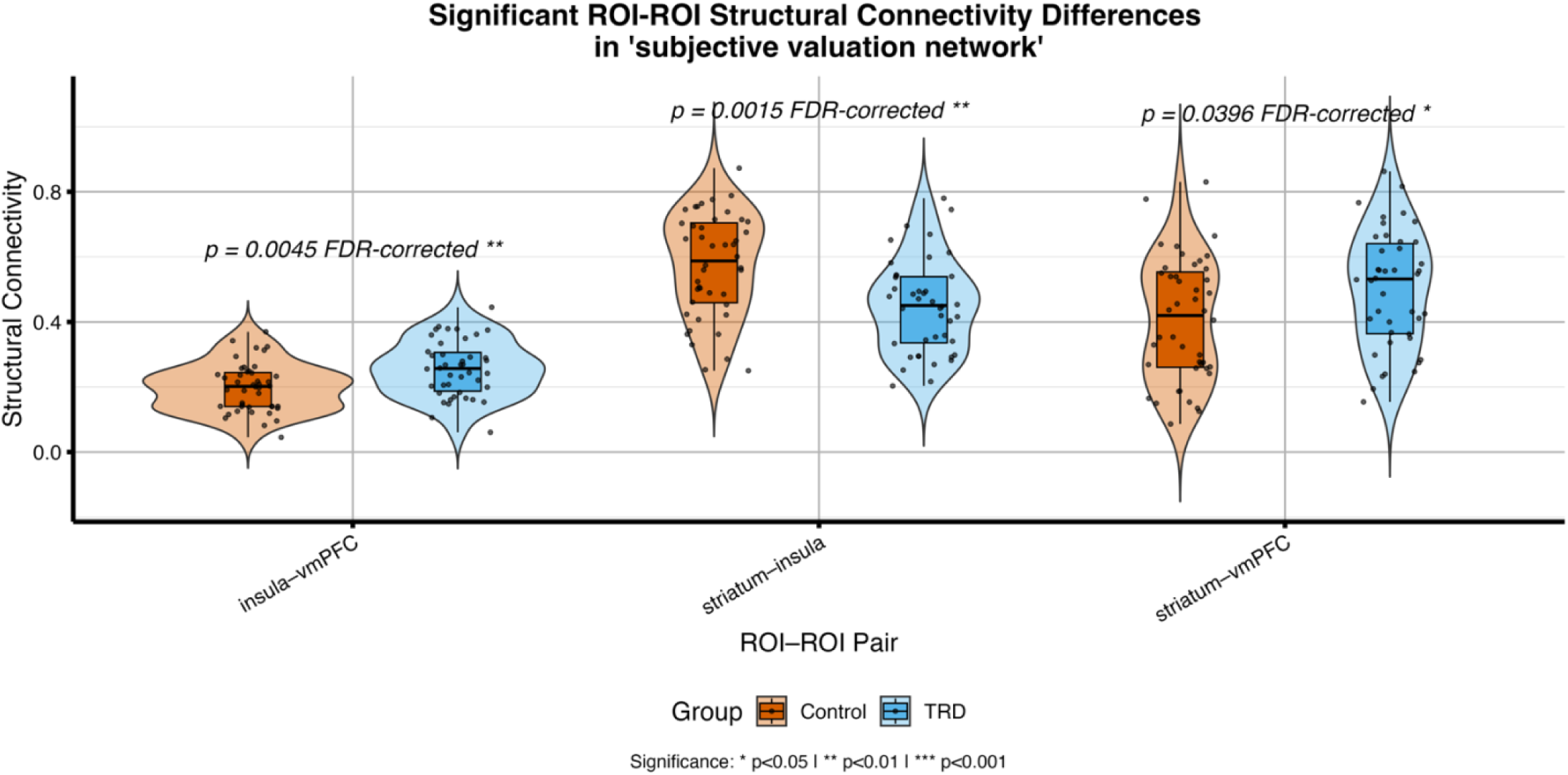
Significantly ROI-ROI structural connectivity differences in the ‘subjective valuation network’ (SVN): Violin plots with overlaid boxplots (median +/− interquartile ranges) and individual data points show group differences for the three key white matter tracts with significantly different structural connectivity in the SVN. From left to right, the insula-vmPFC structural connectivity was increased in TRD versus controls (p = 0.0045, FDR-corrected), striatum-insula structural connectivity was decreased in TRD versus controls (p = 0.0015, FDR-corrected, and striatum-vmPFC structural connectivity was increased in TRD versus controls (0.04), FDR-corrected). Significant annotations placed above each pair; legend indicates group colors.

Correlational analysis of individual ROI-ROI pairs: We examined correlations between white matter tract connectivity and the collected clinical measures (HAMD, PHQ9, MOCA) as well as age, independent of group membership (see figure 5).

**Figure 5:**
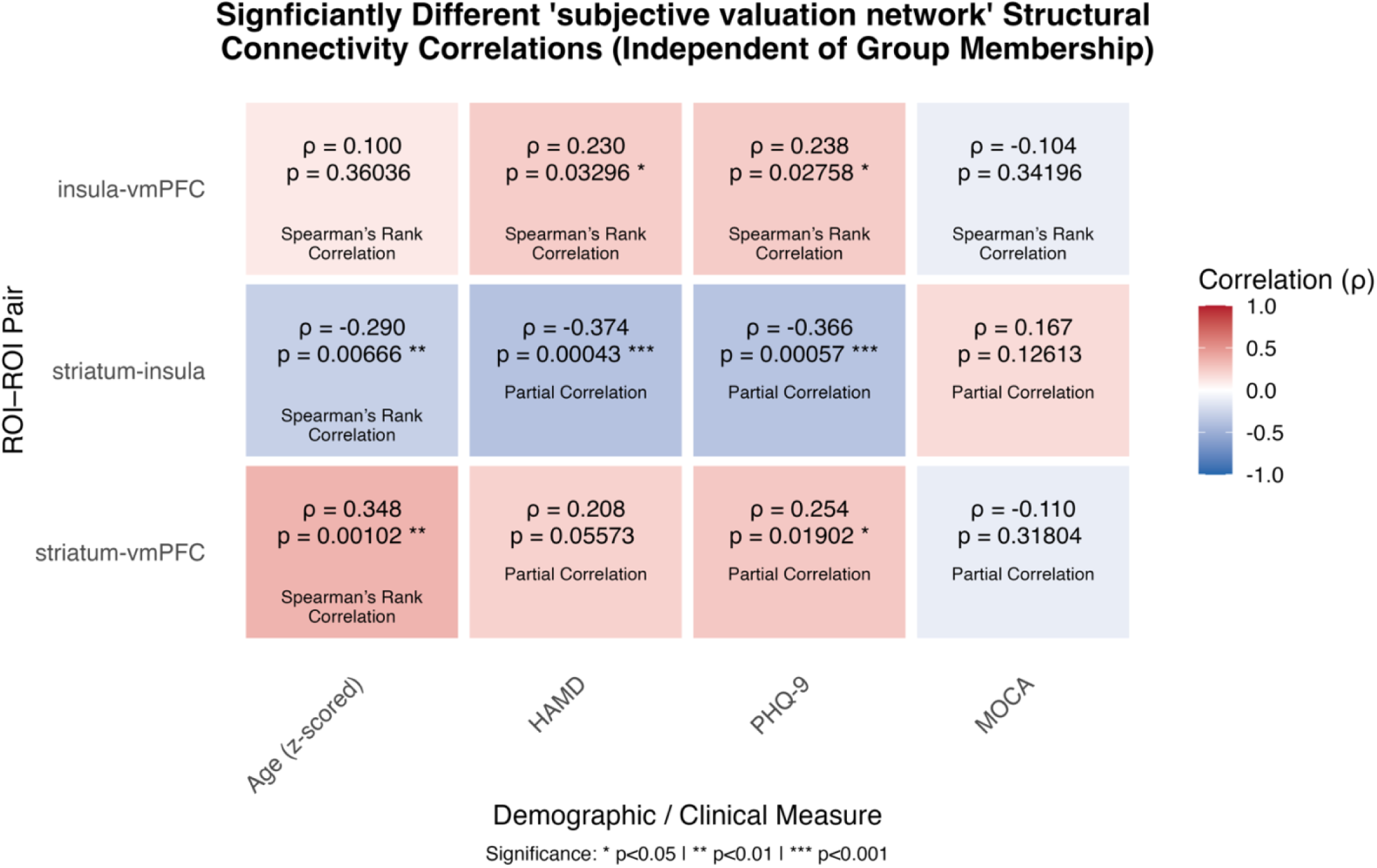
Correlations between structural connectivity of significantly altered ‘subjective valuation network’ (SVN) tracts and demographic/clinical measures (independent of group membership). Each cell reports the correlation coefficient (ρ) and p-value with color indicating the sign and magnitude of the association (red = positive, blue = negative). Insula-vmPFC structural connectivity showed both positive correlation with HAMD (ρ = 0.23, p = 0.33) and PHQ9 (ρ = 0.24, p = 0.028) clinical measures. Striatum-insula structural connectivity demonstrated a significant negative correlation with age (ρ = –0.29, p = 0.0067) and a significant partial correlation (with age as a covariate) with HAMD (ρ = –0.37, p = 0.00043) and PHQ9 (ρ = –0.37, p = 0.00057) clinical measures. Striatum-vmPFC structural connectivity significantly correlated with age (ρ = 0.35, p = 0.001) and significantly partially correlated with PHQ9 (ρ = 0.25, p = 0.02). Significant annotations placed above each pair.

Analysis of the ***insula-vmPFC*** revealed no significant correlation with age (p = 0.36) or MOCA (p=0.34) but showed positive correlation with both HAMD (ρ = 0.23, p = 0.33) and PHQ9 (ρ = 0.24, p = 0.028) clinical measures.

The structural connectivity of the ***striatum-insula*** demonstrated a negative correlation with age (ρ = –0.29, p = 0.0067) and partially negatively correlated (with age regressed out) with HAMD (ρ = –0.37, p = 0.00043) and PHQ9 (ρ = –0.37, p = 0.00057) clinical measures and did not significantly partially associate with MOCA (p = 0.13).

The structural connectivity of the ***striatum-vmPFC*** significantly correlated with age (ρ = 0.35, p = 0.001), did not significantly partially correlate with HAMD (p = 0.056) and MOCA (p = 0.32), but was found to significantly partially correlate with PHQ9 (ρ = 0.25, p = 0.02)

Figure 5 details the complete results of this exploratory correlational analysis offering insight into the directed clinical associations for hypothesis generating purposes.

## Discussion

This study revealed three key findings regarding white matter architecture in TRD: (1) widespread microstructural disruptions in major white matter tracts, particularly affecting interhemispheric and fronto-occipital pathways; (2) selective alterations in structural connectivity within the SVN, despite preserved global network metrics; and (3) specific patterns of altered connectivity in key subcircuits that may reflect both compensatory and maladaptive changes in TRD.

### TBSS

Voxelwise TBSS analysis revealed widespread FA reductions in TRD when compared to controls, consistent with previous findings in MDD and TRD.(18,55) Most pronounced differences in white matter microstructural integrity were located in interhemispheric, frontal-occipital, and frontoparietal pathways, and other long range tracts. In the corpus callosum, FA was lower in TRD participants in both the forceps minor and splenium indicating a disrupted exchange of reward and emotion related signals across hemispheres,(2) additionally reflecting deficits in the integration of contextual visual information during self-referential processing implicated in depression *(Table 2).*(12,55)

Decreased FA in the left IFOF and left superior and inferior longitudinal fasciculi demonstrate weakened ventral-stream and frontoparietal communication, tracts of known networks critical for coupling emotional salience with cognitive control and updating working memory.(12,28,36,40,56) Such white matter disturbances may hamper the switching between the DMN and executive control networks contributing to excessive rumination and motivational anhedonia which are characteristic behavioral features of TRD.(30,32)

The persistent significance of reduced FA in the splenium and forceps minor in age-covariate models suggests these tracts are particularly vulnerable in TRD and likely represents a core pathology specific to the disorder *(Table 3).*(2,27,51) This robustness underscores the role of the corpus callosum in TRD and the importance of interhemispheric communication for cognitive and emotional processing. Collectively, these microstructural deficits in frontolimbic white matter pathways coincide with the reward hyposensitivity, reduced motivation, and anhedonia seen in TRD.(12,28,36,40,56) These findings suggests these disruptions contribute to impaired subjective valuation and reward processing in TRD, further motivating subsequent white matter analyses of the SVN.

## The ‘subjective valuation network’ (SVN)

### Global SVN Network Analyses

In contrast to the widespread TBSS findings revealing reduced FA in several key white matter tracts, macro-level network analyses of the SVN revealed no group level differences; mean FA (t-test, p = 0.78), *composite network connectivity* (Mann-Whitney-U; p = 0.28) (Figure 3). This suggests that while long range white matter structural integrity tracts of the SVN are compromised, the overall regional FA and structural connectivity of the SVN may be preserved, underscoring the value of subcircuit-level analyses that may reveal complex patterns unique to TRD.

However, while mean FA values did not significantly differ between groups, the mean FA of the SVN significantly negatively correlated with age for the TRD group (Age: p = 0.0060 FDR-corrected; (ρ = –0.45) and not the Control group (p = 0.087) suggesting a potential faster rate of decay of white matter integrity in the SVN specific to TRD (Figure 2).

Given the white matter integrity differences broadly observed in TRD and previous findings that MDD is characterized by altered structural connectivity across major white matter pathways,(12) the global network metrics may not fully capture the more localized sub-circuit level white matter pathology in the SVN; further underscoring the importance of examining the white matter structural connectivity of individual ROI-ROI fiber tracts connecting the SVN.

It is important to note that white matter integrity (assessed via FA) and probabilistic connectivity (streamline counts) are related but not identical metrics and capture complementary not identical white-matter properties.(19)

### Individual structural connectivity ROI-ROI pairs within the SVN

Three ROI-ROI pairs’ structural connectivity within the SVN were found to be significantly different between groups: striatum-vmPFC, insula-vmPFC (both connections: TRD > Control), and insula-striatum (Control > TRD) (Figure 4). These findings suggest a complex pattern of structural hyperconnectivity and hypoconnectivity within the SVN subcircuits that may underlie the core symptoms of TRD.

### Increased insula-vmPFC structural connectivity in TRD

Increased structural connectivity between the insula-vmPFC in TRD may indicate a maladaptive neural circuit that amplifies negative self-referential processing and rumination.(2,12,18,25,34,56) The vmPFC is implicated in the self-referential thought, valuation of choices, and top-down emotion regulation,(25,35,56–58) while the insula is critical for integrating interoceptive signals and detecting salience.(25,35,59) Hyperconnectivity between the insula-vmPFC in TRD could lead to pathological amplification of self-referential processes mediated by the vmPFC and integration of internal bodily signals via the insula reflecting a maladaptive hypersynchrony, potentially forming a ‘locked-in loop’ of negative interoceptive salience and valuation, promoting persistent rumination.(18,25,30,34,36,59) This is consistent with findings of over recruitment of the salience network in MDD fostering excessive negative internal focus.(25) In the context of motivated-decision making behavior, it could mean a possible miscalibration of salience attributed to negative or devalued outcomes leading to maladaptive motivational states.(2,12,28–30,35,36) Exploratory correlational analysis independent of group membership revealed a positive association between insula-vmPFC structural connectivity and both HAMD (ρ = 0.23, p = 0.033) and PHQ9 (ρ = 0.24, p = 0.028) depression clinical measures (Figure 5). This supports the idea that increased connectivity in this SVN pathway is linked to greater depression severity further highlighting the potential pathological relevance of this specific SVN subcircuit in TRD.

### Increased striatum-vmPFC structural connectivity in TRD

The observed increase in striatum-vmPFC structural connectivity in TRD could reflect several potential mechanisms. Striatal pathways are classically tied to reward sensitivity and motivational drive(35,38,39,39,40,60) with the vmPFC sending down top-down valuation signals to the striatum.(29,31,38,40,41,61) The increased connection may be a compensatory attempt to overcome functional or neurochemical deficits (e.g. in dopamine signaling) that still contribute to anhedonia.(2,12,29,31,42) Alternatively, it could indicate an aberrant “loop” of negative valuations or a heightened gating mechanism for negative reward predictions.(2,12,27,29,31,35,36,42) It may be that the striatum’s role in learning from reward feedback could be compromised and the hyper-structural-connectivity might paradoxically align with greater negative or uncertain valuations.(2,12,27,29,31,35,36,42) Exploratory correlational analysis independent of group membership showed a significant positive correlation between the structural connectivity of the striatum-vmPFC with age (ρ = 0.35, p = 0.0010) and a significant positive partial correlation with PHQ9 depression scores (ρ = 0.25, p = 0.020) (Figure 5). These associations further support the finding of increased structural connectivity of this SVN pathway in TRD and its potential involvement in maladaptive or compensatory processes ostensibly unique to the disorder.

### Decreased insula-striatum structural connectivity in TRD

Conversely, the detected decreased structural integrity in the insula-striatum seen in TRD suggests a reduced ability to integrate interoceptive and salience signals into reward learning or motivational impetus.(2,27–29,31,36,38,42) The insula typically modulates the motivational salience of incoming reward signals before striatal processing.(35,62) A diminished structural insula-striatum connection may hamper the drive to initiate reward-seeking actions or it may impede the translation of interoceptive reward signals into motivational drive, thereby contributing to behavioral anhedonia.(25,30,31,35,42,43) This aligns with clinical presentations of lowered drive and muted subjective valuation of potential rewards seen in TRD.(2,29,31,36) Exploratory correlational analysis independent of group membership revealed a significant negative correlation with age (ρ = –0.29, p = 0.0067) and significant partial correlations with both HAMD (ρ = –0.37, p = 0.00043) and PHQ9 (ρ = –0.37, p = 0.00057) clinical depression severity scores (Figure 5). This indicates that lower insula-striatum connectivity is associated with greater depression severity, further supporting its role in the anhedonia and motivational deficits observed in TRD.

### Integrating White Matter Findings: Implications for TRD Neuropathology

The pattern of widespread white matter integrity deficits alongside specific contrasting alterations in SVN sub-circuit structural connectivity suggest a complex interplay of factors contributing to TRD pathophysiology. The reduced insula-striatum connectivity coupled with the increased insula-vmPFC-striatum connectivity, may reflect fundamental disruption in salience processing and reward-valuation pathways. Specifically, diminished insula-striatum connectivity could impair the appropriate integration of internal states and external salience with reward processing contributing to anhedonia and reduced motivation. Simultaneously, structural hyperconnectivity in the insula-vmPFC and striatum-vmPFC circuits might facilitate maladaptive loops that amplify negative self-referential thought and rumination potentially by over weighting negative or uncertain outcomes in valuation processes. This aligns with theories of altered salience network and DMN function in depression, in which dysfunctional salience attribution and impaired network switching may sustain internal negative focus and blunt engagement with external rewards.(2,12,17,18,25,27,29,31–34,36,37) The lack of association between these SVN structural connectivity pathways and MOCA scores further supports the notion that these alterations are more specifically related to affective and motivational symptoms rather than general cognitive impairment observed in MDD and TRD,(2,12,13) further evidencing the potential pathophysiological relevance the SVN has with TRD.

### Limitations and Future Directions

Several limitations warrant consideration. The cross-sectional design precludes causal inference with respect to the observed white matter alterations and their relationship to TRD onset or progression; however, it does provide the ability to relate the broad symptomology observed in TRD to underlying pathology. Additionally, medication effects could confound white matter measures. Prior studies have addressed this by controlling for medication type and usage or using medication-free patients. While we observed a potential age-related decline in SVN FA specific to the TRD group, longitudinal studies are needed to rule out potential interventional treatment confounds such as ECT and transcranial-magnetic stimulation treatments as well as illness duration. While the SVN ROI selection was theoretically driven and focused on core network hubs based on converging literature, it excluded potentially relevant regions that warrant investigation, such as the orbitofrontal cortex, amygdala, and hippocampus.(2,12,16–18,25,27–42,51,61) Future studies should employ longitudinal designs, incorporate additional ROIs, and ideally combine structural imaging with functional neuroimaging techniques (e.g. functional-MRI, electroencephalogram, magnetoencephalography, etc.) to better characterize the network dynamics and interplay between structural connectivity with functional and effective connectivity within the SVN. Additionally, building on the growing recognition of psychosocial factors’ critical influence on mental health, future studies should investigate their interplay (e.g. socioeconomic status, early life adversity) with SVN structural connectivity in TRD, as these relationships may help identify additional neurobiological underpinnings.(63) Lastly, the specific sub circuit findings warrant replication in larger cohorts. Investigating how these specific white matter alterations change within the SVN with successful treatment or targeted interventions for anhedonia could also provide valuable insights.

### Conclusion

This study’s multi-level analysis provides compelling evidence that TRD involves distinct patterns of structural connectivity within networks governing subjective reward valuation during motivated decision-making behavior and salience processing. These subcircuit specific alterations, particularly the contrasting differences in insula-striatum, insula-vmPFC, and striatum-vmPFC structural connectivity may represent potential biosignatures of TRD. Together, our findings may offer new targets for therapeutic intervention and biomarker development for this challenging disorder, TRD.

## Acknowledgements

Would like to acknowledge the Kishida Laboratory members for their work and efforts in the experimental design and collection of the data used for this study. This work was supported by the following grant’s awarded to Dr. Kenneth Kishida: National Institute of Health (NIH) R01DA048096, NIH R01MH121099, and NIH R01MH124115; additionally supported by the following grant awarded to Dwayne W. Godwin NIH R01AA016852. This work was also supported by a Neuroscience Clinical Trial and Innovation Center (NCTIC) grant awarded to Wake Forest School of Medicine Clinical and Translational institute.

Benjamin Maas, MS is credited with the scientific hypotheses and conceptualization, original draft preparation, writing – reviewing & editing, data & statistical analyses, methodology, and visualizations. Dwayne W. Godwin, corresponding author is credited with writing – review & editing and further data & statistical analysis support.

Rachel Jones, PhD is credited with the nested ECT original study’s conceptualization, experimental design, methodology, and data collection and curation.

Christina Johnson, PhD and Angela Jiang BS are additionally credited with data collection and curation of the nested original ECT study.

## Disclosures

Benjamin Maas reports no biomedical financial interest or potential conflicts of interests. Rachel Jones, PhD reports no biomedical financial interest or potential conflicts of interests. Christina Johnson, PhD reports no biomedical financial interest or potential conflicts of interest. Angela Jiang reports no biomedical financial interest or potential conflicts of interests.

Dwayne Godwin, PhD reports no biomedical financial interest or potential conflicts of interest.

## Notes

### Competing Interest Statement

The authors have declared no competing interest.

