## Supplementary Materials and Methods for "Complex Patterns of Altered White Matter Structural Connectivity within a ‘subjective valuation network’ in Treatment-Resistant Depression"

### Supplementary Material:

### Supplementary Figures:

| Connectivity Matrix: 'subjective valuation network' (SVN) | | | | |
|---|---|---|---|---|
| ROI in SVN [Row, Column: i, j] | anterior cingulate cortex (ACC) | insula | striatum | ventromedial prefrontal cortex (vmPFC) |
| anterior cingulate cortex (ACC) | Self-Connection (0) | ACC → insula | ACC → striatum | ACC → vmPFC |
| insula | insula → ACC | Self-Connection (0) | insula → striatum | insula → vmPFC |
| striatum | striatum → ACC | striatum → insula | Self-Connection (0) | striatum → vmPFC |
| ventromedial prefrontal cortex (vmPFC) | vmPFC → ACC | vmPFC → insula | vmPFC → striatum | Self-Connection (0) |

| Waypoint Vector: Total Streamline Count Sent Out Per Region of Interest (Row *i* of Connectivity Matrix) |
|---|
| anterior cingulate cortex (ACC) |
| insula |
| striatum |
| ventromedial prefrontal cortex (vmPFC) |

Equations utilized to calculate forward, reverse, and structural connectivity values:

$$Supplemntary\ eqn. (1)\ Normalized\ Connectivity\ Matrix[i, j] = \frac{Connectivity\ Matrix: [i, j]}{Waytotal[i]}$$

$$Supplemntary\ eqn. (2)\ Forward\ Connectivity\ (ROI\ i\ \rightarrow j) = Normalized\ Connectivity\ Matrix[i, j]$$

$$Supplemntary\ eqn\ (3)\ Reverse\ Connectivity\ (ROI\ j\ \rightarrow i) = Normalized\ Connectivity\ Matrix[j, i]$$

$$eqn\ (4)\ Structural\ Connectiviy = Average\ of\ forward\ and\ reverse = \frac{Forward\ Connectivity + Reverse\ Connectivity}{2}$$

Figure S1: Connectivity Matrix and Waypoint Vector for the 'subjective valuation network' and equations utilized to calculate connectivity values: The upper panel shows the 4x4 connectivity matrix derived from probabilistic tractography with each cell [i,j] giving the number of streamlines connecting seed region *i (rows)* to target region *j (columns)*. Diagonal entries represent within-region ("self") connections (with 0 streamline counts). The lower panel displays the 4x1 waypoint vector, listing the total number of streamlines initiated from each ROI (i.e. the row sums of the raw connectivity matrix). These waytotal values were then used to normalize each matrix entry and to compute forward, reverse, and then structural connectivity indices for each ROI-ROI pair using the outlined supplementary equations retrieved from FSL guidelines detailed at https://open.win.ox.ac.uk/pages/fslcourse/lectures/Tract_E4.pdf. Supplementary equations 1-3 were adapted into python code to batch process the assessed connectivity values.

**Supplementary Methods:**

**Diffusion Weighted Imaging (DWI) Preprocessing and TBSS Cluster Labeling Pipeline:**

**DWI Preprocessing (per subject):**

Native scanner mages were converted from DICOM to NIFTI using MRIcron's tool *dcm2niix* (v1.0.20241211). All subsequent image preprocessing and DWI analysis steps were performed using FSL (v6.0.7.4).(48) The following commands represent the general workflow. Subject-level steps were applied iteratively for each participant.

1. Topup (distortion correction):
    a. topup --imain=<acq_pairs> --datain=<acq_params.txt> --out=<topup_results>
2. Brain Extraction (removal of skull and extraneous non-brain matter from non-diffusion weighted image creating a mask subsequently used for *eddy* and *dtifit*):
    a. bet <b0_image.nii.gz> <b0_brain.nii.gz> -m
3. Eddy Current and Motion Correction:
    a. eddy --imain=<data.nii.gz> --mask=<b0_brain_mask.nii.gz> \
    --acqp=<acq_params.txt> --index=<index.txt> \
    --bvecs=<bvecs> --bvals=<bvals> --topup=<topup_results> \
    --out=<eddy_corrected_data>
4. Diffusion Tensor Imaging Model Fitting:
    a. dtifit -k <eddy_corrected_data.nii.gz> -o <dtifit_output_prefix> \
    -m <b0_brain_mask.nii.gz> -r <bvecs> -b <bvals>

**TBSS Cluster Identification and Labeling (Group-Level Results)**:

1. Cluster extraction:
    a. cluster -i <tbss_tfce_corrp_tstatContrastX.nii.gz> -t 0.95 --scalarname="1-p" \
    -oindex=<cluster_index.nii.gz> --osize=<cluster_size.nii.gz> \
    --mm -o <ContrastX-cluster_output> > <ContrastX-cluster_table.txt>
2. Atlas-based white matter tract labeling
    a. atlasquery -a "JHU White-Matter Tractography Atlas" -m <ContrastX-cluster_output.nii.gz>

**BedpostX Model Fitting and Probabilistic Tractography (per subject):**

For each subject FSL's *bedpostx* was run on the preprocessed diffusion data to estimate diffusion parameters for probabilistic tractography.(22,23)

1. Bedpostx (estimate diffusion parameters for probabilistic tractography ran on preprocessed diffusion data):
    a. Bedpostx <SubjectX_diffusion_directory>
2. Inverse Warp files from MNI to Diffusion space (created from TBSS normalization) for ProbtrackX:
    a. invwarp –warp <SubjectX_FA_to_target_warp.nii.gz> --ref <SubjectX_b0.nii.gz> --out <SubjectX_invwarp.nii.gz>
3. ProbtrackX (run in network mode to compute probabilistic tractography across the four ROI masks defining the 'subjective valuation network' (SVN). For each subject:
    a. probtrackx2 \

```
--network \
-x <MASKS_FILE> \
-l \
--onewaycondition \
-c 0.2 \
-S 2000 \
--steplength=0.5 \
-P 1000 \
--fibthresh=0.01 \
--distthresh=0.0 \
--sampvox=0.0 \
--xfm=<SubjectX_xfm_path> \
--invxfm=<SubjectX_invxfm_path> \
--forcedir \
--opd \
-s <SubjectX_bedpostx_dir>/merged \
-m <SubjectX_bedpostx_dir>/nodif_brain_mask \
--dir=<SubjectX_output_dir>
```

**Mean network FA value of the SVN (per subject)**

1. Threshold and binarize top 10% streamline counts of resulting probtrackX image of each subject's SVN probabilistic tractography map:
   a. fslmaths <SubjectX_SVN_map> -thrP 0.9 -bin <SubjectX_SVN_map-top10bin>
2. Multiply thresholded and binarized SVN map by each subject's FA image in MNI standard 1 $mm^3$ space (obtained from TBSS alignment) producing map of FA values of the SVN for each subject:
   a. fslmaths <SubjectX_FA_to_target.nii.gz> -mul <SubjectX_SVN_map-top10bin> <SubjectX_FA_SVN>
3. Output mean network FA value of the SVN (non-zero value of the SVN image with a minimum of 0.15 value). For each subject.
   a. fslstats <SubjectX_FA_SVN> -l 0.15 -m
